# How Alzheimer’s Aβ propagates and triggers tau pathology in intact neurons

**DOI:** 10.1101/2024.08.28.608712

**Authors:** Tomoya Sasahara, Hongo Shoko, Mika Ito, Naomi Takino, Shin-ichi Muramatsu, Akiyoshi Kakita, Minako Hoshi

**Author notes:** Correspondence and requests for materials should be addressed to M.H.

## Abstract

Alzheimer’s disease begins with Aβ accumulation, progresses to tau aggregates and results in widespread neurodegeneration^1-3^. Simultaneous propagation of Aβ aggregates from very limited to wide and distant brain regions is one of the outstanding events in the early stage. Here, we demonstrate that the neurovascular unit^4,5^ comprising capillaries and pericytes, the machinery supplying oxygen and glucose to neurons, is also the machinery for propagating effector molecules that impose Alzheimer’s pathologies on intact neurons. We discovered two distinct signaling cascades, one activated in capillary endothelial cells and the other in pericytes. At the origin of either cascade, we identified amylospheroid (ASPD)^6,7^, a highly toxic 30-mer assembly of Aβ, and its sole target NKAα3^8^, a neuron-specific isoform of Na^+^,K^+^-ATPase^9,10^ but present in endothelial cells and pericytes. In endothelial cells, ASPD binding to NKAα3 releases angiotensin II, which increases β-secretase in intact neurons, causing a huge increase of Aβ_42_and resultant accumulation. In pericytes, ASPD binding to NKAα3 releases an unknown effector molecule that activates δ-secretase in intact neurons, further augmenting Aβ_42_and producing pathogenic tau_1-368_ fragment. Thus, Aβ and tau pathologies are directly linked. Stopping the signaling cascades near the origin by inhibiting ASPD-NKAα3 interaction^8^ may provide a new therapeutic approach.

## Main

Alzheimer’s disease (AD) is a progressive dementia, accounting for ∼60% of dementia cases. It begins with the accumulation of aggregated forms of amyloid-β protein (Aβ) in limited neocortical regions and progresses to accumulate cleaved and hyperphosphorylated forms of tau protein^1-3^. Both Aβ and tau aggregates spread throughout the brain, eventually leading to neurodegeneration^2,3^. It is of critical importance for combating AD at its earliest stage to understand the molecular mechanisms underlying the accumulation and spreading of Aβ aggregates, the events that occur in the earliest stage of the disease. However, our understanding of these processes is still very limited.

Aβ is a protein of mostly 38-42-residues produced inside neurons by sequential cleavages of amyloid precursor protein (APP) with β- and γ-secretases^11^. Aβ is constantly produced in all neurons in all people, as it plays vital roles in maintaining synaptic function and memory^12^. Normally, Aβ is cleared from the brain without accumulation, but under certain conditions, Aβ forms toxic aggregates that exert various effects on neurons depending on the forms and sizes of the aggregates, from synaptic dysfunction to neurodegeneration^8,13^.

We have isolated amylospheroid (ASPD), ∼30-mer assembly (∼128 kDa) of Aβ, as a potent neurotoxic assembly from human patient brains^6,7^. Its sole target is a Na^+^ pump, NKAα3 (neuron-specific isoform of Na^+^,K^+^-ATPase)^8^. ASPD binds to the extracellular region of NKAα3 with a high affinity (8 nM) and suppresses ATPase activity^8^.

Although Aβ is produced in every neuron, ASPD is formed only inside excitatory neurons and secreted from the synaptic terminal^14^. Secreted ASPD damages NKAα3-expressing inhibitory neurons^15^ that project to the excitatory neurons, thereby causing local damage that destroys both types of neurons^14^. ASPD begins accumulating, most likely in limited neocortical regions, long before behavioral deficits become apparent, as demonstrated with 5xFAD mice^16^. ASPD increases in parallel with the disease severity, eventually reaching ∼60% of all nonfibrillar Aβ aggregates in the neocortex of patients with AD^8^. Fibrillar aggregates of Aβ also accumulate and spread throughout the brain but subsequently to ASPD^16^.

Thus, ASPD formation in a limited number of excitatory neurons appears to be the earliest event in AD. How then ASPD or Aβ pathology spreads over a wide brain area is a question we would like to address here. ASPD certainly damages projected neurons, but no direct transport of ASPD beyond a synapse has been detected. For spreading, it would be more efficient than direct transport of ASPD to propagate “conditions”, such as effector molecules that induce aggregation of Aβ, as all neurons produce Aβ. In the early stages of AD, Aβ simultaneously accumulates in distant medial frontoparietal regions^17,18^. A recent transcriptomic analysis of two cohorts has found that a set of genes related to Aβ pathology is predominantly expressed in these regions, conferring susceptibility to Aβ pathology^19^.

However, Aβ does not accumulate under physiological conditions, even in these regions. A “condition” that triggers the accumulation of Aβ must be induced simultaneously across the wide regions.

In this regard, we should be aware that every neuron stays proximate to a blood capillary (Extended Data Fig. 1)^20^, in which endothelial cells and pericytes communicate via effector molecules. Neurons consume huge amounts of energy, requiring continuous supply from blood. The brain requires 50-60 mL/min of blood per 100 g brain to work properly^20^. That is why neurons and capillaries form the “neurovascular unit” (NVU)^4,5^. Because blood capillaries are connected and spread across the entire brain, they could act as a conduit for effector molecules that impose Aβ pathology on intact neurons. It is well known from studies on human autopsy brains that vascular disorders worsen Aβ pathologies^21^, although it remains obscure whether there is a direct link between vascular damage and the spreading of Aβ pathology. Of particular interest here is that NKAα3, the sole target of ASPD^8^, though widely described as a neuron-specific isoform^9,10^, is expressed in endothelial cells of brain capillaries^22^. It is then plausible that ASPD damages endothelial functions. Indeed, ASPD treatment of blood vessels inhibits vasorelaxation, indicating that a pathological signaling cascade is activated through NKAα3^22^. Such a mechanism might also be effective in propagating Aβ pathology. We, therefore, constructed a new 3D culture system mimicking the NVU, consisting of neurons, endothelial cells and pericytes, and examined the hypothesis.

Experiments with the new culture system unequivocally indicated that Aβ pathology propagates through the NVU by spreading the “conditions” (i.e. effector molecules) that cause a huge increase in the production of Aβ_42_ and their resultant accumulation. We demonstrate that two pathological signaling cascades, distinct between endothelial cells and pericytes, are activated by ASPD-NKAα3 interaction. An unexpected finding here is that the pathological cascade in pericytes activates δ-secretase that initiates tau pathology. Aβ pathology and tau pathology are directly linked.

### Blood ASPD increases neuronal APP cleavage

Capillaries comprise inner endothelial cells and the outer basement membrane, in which pericytes are embedded and interact directly with endothelial cells (Fig. 1a left). Neurons stretch their endings near capillaries but do not contact them directly. To mimic this arrangement, we divided the culture well into upper and lower sections with a membrane insert, on which endothelial cells and pericytes, both from human brain capillaries, were grown (Fig. 1a right). This setup allows only endothelial cells to be exposed to the upper medium (U-med) corresponding to blood, while pericytes stay in touch with endothelial cells. Mature neocortical neurons were seeded in the bottom to avoid direct contact with either endothelial cells or pericytes (Fig. 1a right). Most (79.1%) ASPD added to U-med at 36.8 nM remained there after 48 h. Only a trace amount (0.11 nM at maximum) appeared in the lower medium (L-med) (Extended Data Fig. 2). This amount of ASPD would have little effect on neurons in the bottom of the well, as ASPD has an EC_50_ value of 18 ± 2.4 nM for neurotoxicity and *K*_D_ value of 7.8 ± 2.5 nM for NKAα3^8^. We used this NVU mimicking system for further study.

**Fig. 1.**
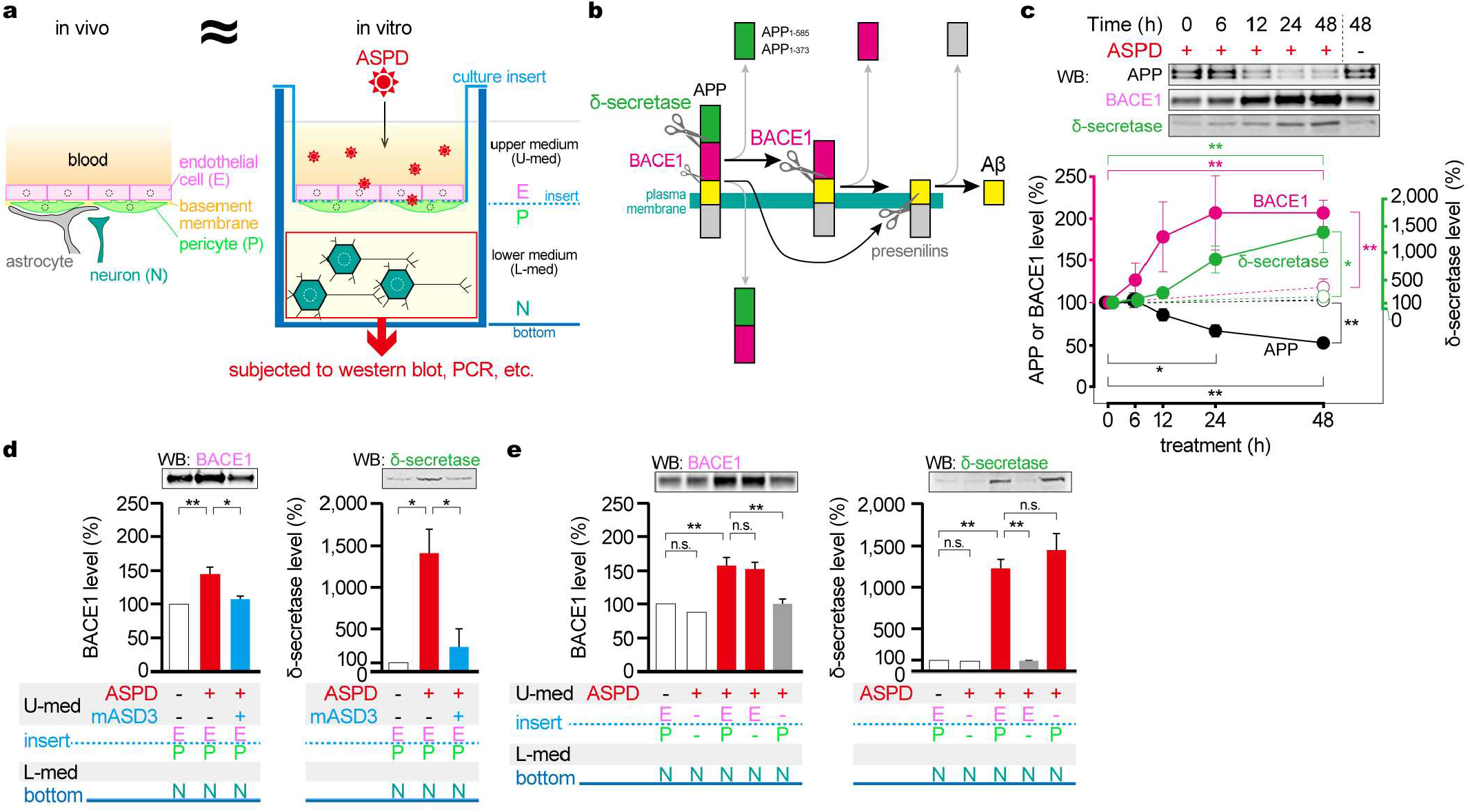
Effects of ASPD on the new culture system mimicking the neurovascular unit (NVU). **a**, Spatial arrangement of neurons (N), endothelial cells (E) and pericytes (P) in NVU (left) and in the 3D culture system mimicking the NVU (right). **b**, Two pathways for the production of Aβ. The one initiated by δ-secretase allows more efficient cleavage of APP^23^. **c**, Quantification by western blotting (*n* = 4) of APP (double band due to variations in glycosylation), BACE1 and δ-secretase^23^ in neuronal extracts after the addition of ASPD (34.1 nM) to U-med. **d**, Quantification (*n* = 3) of BACE1 and δ-secretase after 48-h addition of ASPD (56.4 nM) to U-med with or without pretreatment of ASPD-specific mASD3 antibody^7^. **e**, Effects of ASPD treatment (48 h, 35.4 nM) on the four combinations of cocultures with neurons, as measured by BACE1 and δ-secretase levels (*n* = 4). In **c-e**, data represent the mean ± S.E. normalized to actin levels. \**p* < 0.05, \*\**p* < 0.01, n.s. *p* > 0.05 (one-way ANOVA with Scheffé’s method). Compositions of the culture system (E, endothelial cell; P, pericyte; N, neuron) and medium conditions (inclusion of ASPD, etc.) are shown below each bar graph. For the raw data of **c, d** and **e**, refer to Extended Data Fig. 3**a, b** and **d**, respectively.

We first examined changes in the amounts of APP and of three key enzymes for Aβ production in neurons, namely, β-, δ- and γ-secretases (Fig. 1b), by western blotting. β-secretase is also called BACE1 (β-site APP cleaving enzyme). δ-secretase is a 46-kDa asparagine endopeptidase produced by self-cleavage of the 56-kDa inactive proenzyme^23^. It is a widely distributed lysosomal protease, acting usually as a scavenger. These two enzymes are rate limiting in Aβ production, whereas the catalytic subunit of the third enzyme, γ-secretase, or presenilin-1, is not^24,25^. The addition of ASPD (34.1 nM) to U-med slowly decreased APP levels in neurons to a statistically significant level after 24 h and to 53% after 48 h (Fig. 1c). The two rate-limiting enzymes increased in parallel with the decrease in APP: BASE1 two-fold, whereas δ-secretase fifteen-fold (Fig. 1c). These changes are ASPD-dependent (open circles in Fig. 1c).

When the binding of ASPD to NKAα3 was blocked by mASD3^7,8,22^, an ASPD-specific monoclonal antibody, the protein levels of BACE1 and δ-secretase remained the same (Fig. 1d). The protein level of presenilin-1 also increased, but independent of ASPD (Extended Data Fig. 3c). Thus, ASPD binding to NKAα3 in endothelial cells and/or pericytes increases BACE1 and δ-secretase in neurons, but not presenilin-1.

Next, to identify the cells responsible for these increases, we prepared four combinations of cocultures with neurons and measured the protein levels in the neurons 48 h after ASPD addition (Fig. 1e). BACE1 increased by the ASPD treatment of endothelial cells alone. δ-secretase was activated by the ASPD treatment of pericytes alone. No increase was detected if neither of them was present. It is thus evident that endothelial cells and pericytes act independently on neurons.

### Angiotensin II induces Aβ accumulation via BACE1

As endothelial cells and neurons do not contact directly, we presumed ASPD acts on endothelial cells to release an effector molecule that increases BACE1 in neurons. As a candidate, we examined angiotensin II (Ang II), a peptide hormone that regulates vasoconstriction^26^. Ang II is produced by angiotensin-converting enzyme (ACE) and binds to Ang II type 1 receptor (AT1R) or type 2 receptor (AT2R) for its function^26^. The addition of ASPD indeed increased Ang II in U-med nearly twenty-fold (from 1.8 ± 0.4 to 33.4 ± 0.8 nM (*n* = 4)) after 6 h. At that time, BACE1 began to increase in neurons (see Fig. 1c). Either blocking Ang II production in endothelial cells or inhibiting the action of AT2R in neurons almost abolished the increase of BACE1 in neurons after 48 h of ASPD exposure (Fig. 2a). For the blocking, we used 0.1 µM captopril, an ACE-specific inhibitor, or 1 µM PD123319, an inhibitor specific to AT2R. Treating neurons with Ang II (100 nM) increased BACE1, but not if AT2R action was blocked (Extended Data Fig. 4b). Conversely, the activation of AT2R with a subtype-selective agonist (100 nM CGP42112) increased BACE1 (Extended Data Fig. 4c). These findings indicate that Ang II, produced by ACE in endothelial cells, induces BACE1 in neurons through AT2R.

**Fig. 2.**
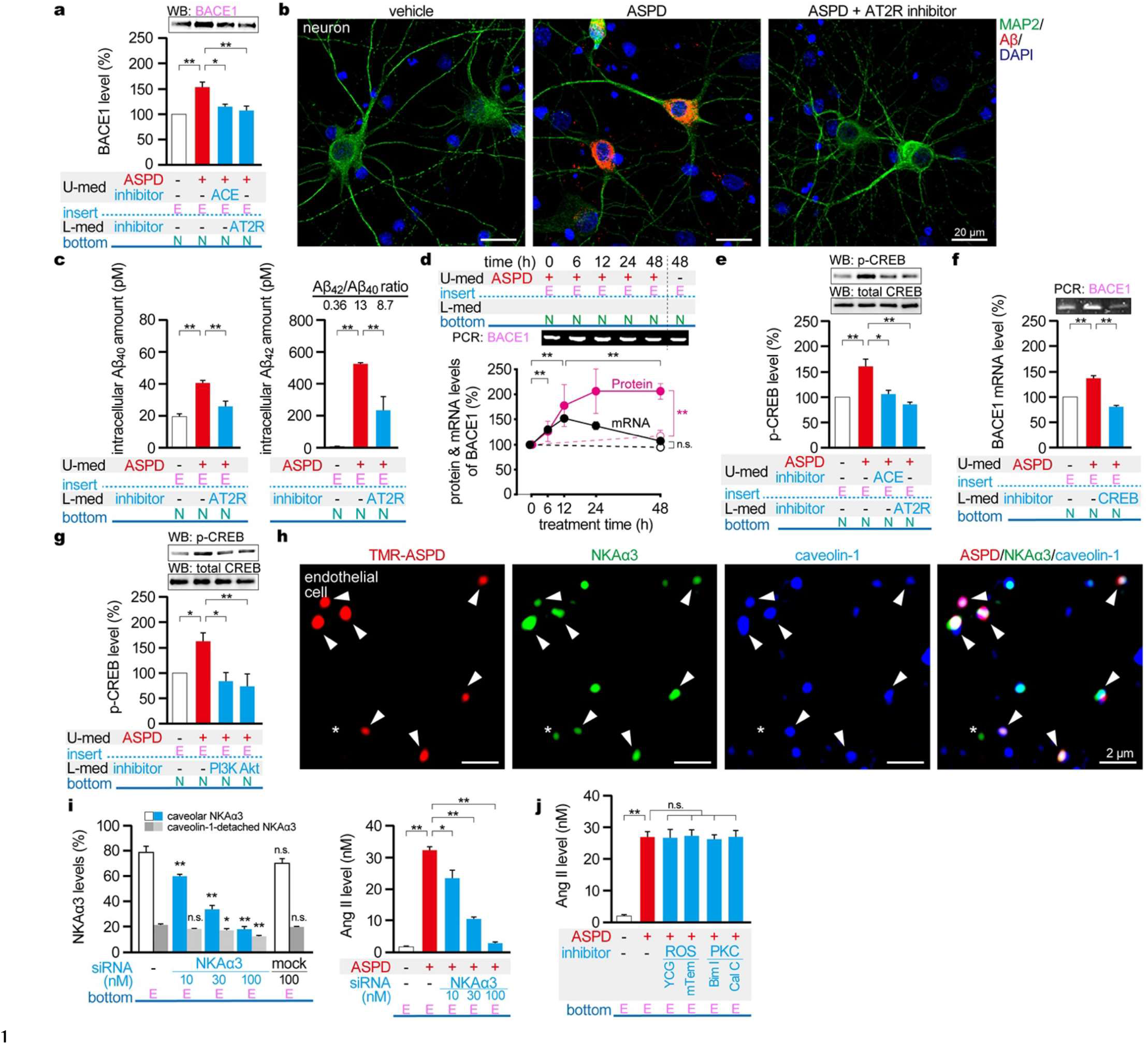
Release of Ang II by ASPD treatment of capillary endothelial cells and resultant Aβ accumulation in neurons. **a**, BACE1 increase in neurons, normalized to actin, after ASPD treatment (48 h, 39.8 nM) of endothelial cells (*n* = 4). Inhibitors of ACE (0.1 µM captopril) and AT2R (1 µM PD123319) suppressed the increase. **b and c**, Aβ increase in neurons expressing human APP Swedish mutation detected by immunostaining (**b**) or Aβ ELISA (**c**) after ASPD treatment (**b**, 60.0 nM; **c**, 25.6 nM) of endothelial cells for 48 h. In **b**, Aβ was detected by immunostaining with an N-terminal-end-specific antibody recognizing human Aβ but not APP. In **c**, the levels of Aβ_40_ and Aβ_42_ in the neuronal extracts were determined by ELISA specific to each Aβ (*n* = 3). **d**, Time-dependent changes in *BACE1* mRNA in neurons determined by semi-quantitative RT-PCR after ASPD treatment (57.8 nM) of endothelial cells (*n* = 4; normalized to *GAPDH* mRNA). For comparison, changes in BACE1 protein in Fig. 1c are shown. **e**, p-CREB increase in neurons determined by western blotting after ASPD treatment (12 h, 39.8 nM) of endothelial cells (*n* = 3; normalized to total CREB). **f**, Inhibition of *BACE1* mRNA increase by treatment of neurons with 10 µM 666-15 (a CREB inhibitor) (*n* = 4; normalized to *GAPDH* mRNA). *BACE1* mRNA was elevated by ASPD treatment (12 h, 38.9 nM) of endothelial cells. **g**, Inhibition of the p-CREB increase by treatment of neurons with a PI3K inhibitor (1 µM wortmannin) or Akt inhibitor (10 µM Akt inhibitor I). p-CREB was increased by ASPD treatment (12 h, 52.4 nM) of endothelial cells (*n* = 4). **h**, Immunostaining (NKAα3, caveolin-1) of endothelial cells after 10 min treatment with tetramethyl rhodamine-labeled ASPD (TMR-ASPD; 32.9 nM). TMR-ASPD colocalized with NKAα3 associated with caveolin-1 (white arrowheads) but not with NKAα3 detached from caveolin-1 (asterisks). **i**, Effects of pretreatment of endothelial cells with NKAα3 siRNA on levels of two types of NKAα3 (caveolar and the caveolin-1-detached; left; *n* = 10) and Ang II (right; *n* = 4) after ASPD treatment (6 h, 48.2 nM) of endothelial cells. Numbers of each type of NKAα3 per cell area were determined by immunostaining, and the relative values are shown; the sum of the two types of NKAα3 in siRNA-untreated cells was taken as 100%. **j**, Quantification of Ang II as in **i** (*n* = 4) after ASPD treatment (6 h, 23.5 nM) of endothelial cells, with or without an inhibitor for ROS (50 µM YCG-063 (YCG) and 100 µM mito-tempol (mTem)) or for PKC (5 µM bisindolylmaleimide I (Bim I) and 0.3 µM calphostin C (Cal C)). In **a, c-g** and **i-j**, data represent the mean ± S.E. \**p* < 0.05, \*\**p* < 0.01, n.s. *p* > 0.05 (one-way ANOVA with Scheffé’s method). For the raw data of **a, b, d, e, f** and **g**, refer to Extended Data Fig. 4**a**, 4**d**, 5**a**, 5**b**, 5**c** and 5**d**, respectively.

The observed increase of BACE1 would promote Aβ production in neurons, as BACE1 is a rate-limiting protease for Aβ production^11,24^. This possibility was examined using mature neocortical neurons expressing human APP carrying the Swedish mutation (APP Swedish mutation), one of the known genetic mutations in familial AD. After treating endothelial cells for 48 h with ASPD, Aβ accumulated predominantly in the cell bodies of the neurons (Fig. 2b). This accumulation entirely depended on ASPD treatment and was abolished by the inhibitor specific to AT2R (Fig. 2b). Thus, ASPD treatment of endothelial cells results in accumulation of Aβ in neurons because BACE1 increases via the ACE/Ang II/AT2R cascade.

The Swedish mutation in APP elevates the production of both Aβ_1-40_ and Aβ_1-42_ by ∼6-fold^27,28^. It was therefore surprising that the 48-h ASPD treatment of endothelial cells elevated Aβ production even further: Aβ_40_ two-fold and Aβ_42_ as much as ∼70-fold (Fig. 2c). The AT2R-specific inhibitor eliminated the increase of Aβ_40_ but blocked only ∼2/3 of the Aβ_42_ increase (Fig. 2c right). The reason for this observation is unclear.

Next, we explored how Ang II increases BACE1 protein in neurons. As *BACE1* mRNA significantly increased several hours ahead of BACE1 protein (Fig. 2d), transcription of the *BACE1* gene must have been up-regulated. As a transcriptional factor, we examined CREB, the cAMP-responsive element binding protein. Once activated by phosphorylation at Ser133 (p-CREB), this protein promotes *BACE1* transcription by binding to the CRE site of its promoter^29^. Treating endothelial cells with ASPD increased p-CREB by ∼60% after 12 h with no change in the total amount of CREB (Fig. 2e). At this time, the amount of *BACE1* mRNA in neurons elevated by ∼50%, reaching the maximum (Fig. 2d). Experiments with ACE- or AT2R-specific inhibitors showed that the increase in p-CREB depends on Ang II (Fig. 2e). Furthermore, ASPD treatment of endothelial cells caused no increase in *BACE1* mRNA if a CREB-specific inhibitor (10 µM 666-15)^30^ was present (Fig. 2f). These observations indicate that, in neurons, the ACE/Ang II/AT2R cascade leads to the phosphorylation of CREB at Ser133, which promotes the transcription of *BACE1* gene to increase BACE1 protein.

We then examined if the phosphatidylinositol-3 kinase (PI3K)/Akt cascade^31^ links AT2R and p-CREB^32^. The treatment of neurons with a specific inhibitor of PI3K (1 µM wortmannin) or of Akt (10 µM Akt inhibitor I) entirely blocked the increase of p-CREB after 12-h of ASPD treatment in endothelial cells (Fig. 2g). These two inhibitors specific to the PI3K/Akt cascade also suppressed the increase of p-CREB induced by the 12-h treatment of 100 nM Ang II in neurons (Extended Data Fig. 5e). These data indicate that, in neurons, the PI3K/Akt cascade works downstream of AT2R and leads to p-CREB.

As mentioned earlier, the sole target of ASPD is NKAα3, whether in neurons^8^ or endothelial cells^22^. In neurons, caveolin-1 is hardly expressed (Extended Data Fig. 5f), and no signal was detected for the coexistence of NKAα3 and caveolin-1 (Extended Data Fig. 5g). However, in endothelial cells, in which caveolin-1 is abundant (Extended Data Fig. 5f), three quarters (74.9 ± 11.3% (*n* = 4)) of NKAα3 coexisted with caveolin-1 (Fig. 2h and i left). Indeed, super-resolution microscopy showed that, of the ASPD signals coexisting with NKAα3, 91.1 ± 1.6% (*n* = 4) colocalized with caveolin-1 in endothelial cells (arrowheads in Fig. 2h). We then tested if caveolar NKAα3 is responsible for Ang II release by knocking down NKAα3 with small interfering RNA (siRNA). Surprisingly, the knockdown suppressed caveolar NKAα3 predominantly (Fig. 2i left), decreasing the release of Ang II in strict parallel (Fig. 2i right). Thus, it is unequivocal that caveolar NKAα3 mediates ASPD action leading to Ang II release.

We previously showed that ASPD binding to caveolar NKAα3 in endothelial cells produces reactive oxygen species (ROS), which activate protein kinase C (PKC), eventually leading to the inactivation of endothelial nitric oxide synthase (eNOS) by PKC-mediated phosphorylation^22^. Here, we investigated whether or not the ROS/PKC/eNOS inhibition cascade is involved in Ang II release downstream of the interaction between ASPD and caveolar NKAα3 and found no inhibitors against ROS or PKC caused a change in Ang II release after ASPD treatment (Fig. 2j). Accordingly, ASPD activates at least two signaling cascades. The ROS/PKC/eNOS inhibition cascade leads to reduced blood flow by inhibiting vasoconstriction^22^. The ACE/Ang II release cascade induces BACE1 expression in healthy neurons and causes a huge increase in the production of Aβ_42_, leading to their accumulation. Thus, ASPD damages blood vessels and propagates Aβ pathology to healthy neurons through its binding to caveolar NKAα3 in endothelial cells.

### ASPD triggers tau pathology via pericytes

Pericytes express NKAα1, the ubiquitous isoform of Na^+^,K^+^-ATPase, to which ASPD does not bind (Extended Data Fig. 6a)^8^. However, it was unknown whether pericytes also express NKAα3 before our study. To our surprise, NKAα3 is abundant in pericytes (Fig. 3a), as high as in neurons (Extended Data Fig. 6b). In contrast, pericytes express caveolin-1 hardly any (∼1/30 in endothelial cells) (Extended Data Fig. 5f), consistent with the lack of colocalization signal with NKAα3 (Fig. 3a), like neurons (Extended Data Fig. 5g).

**Fig. 3.**
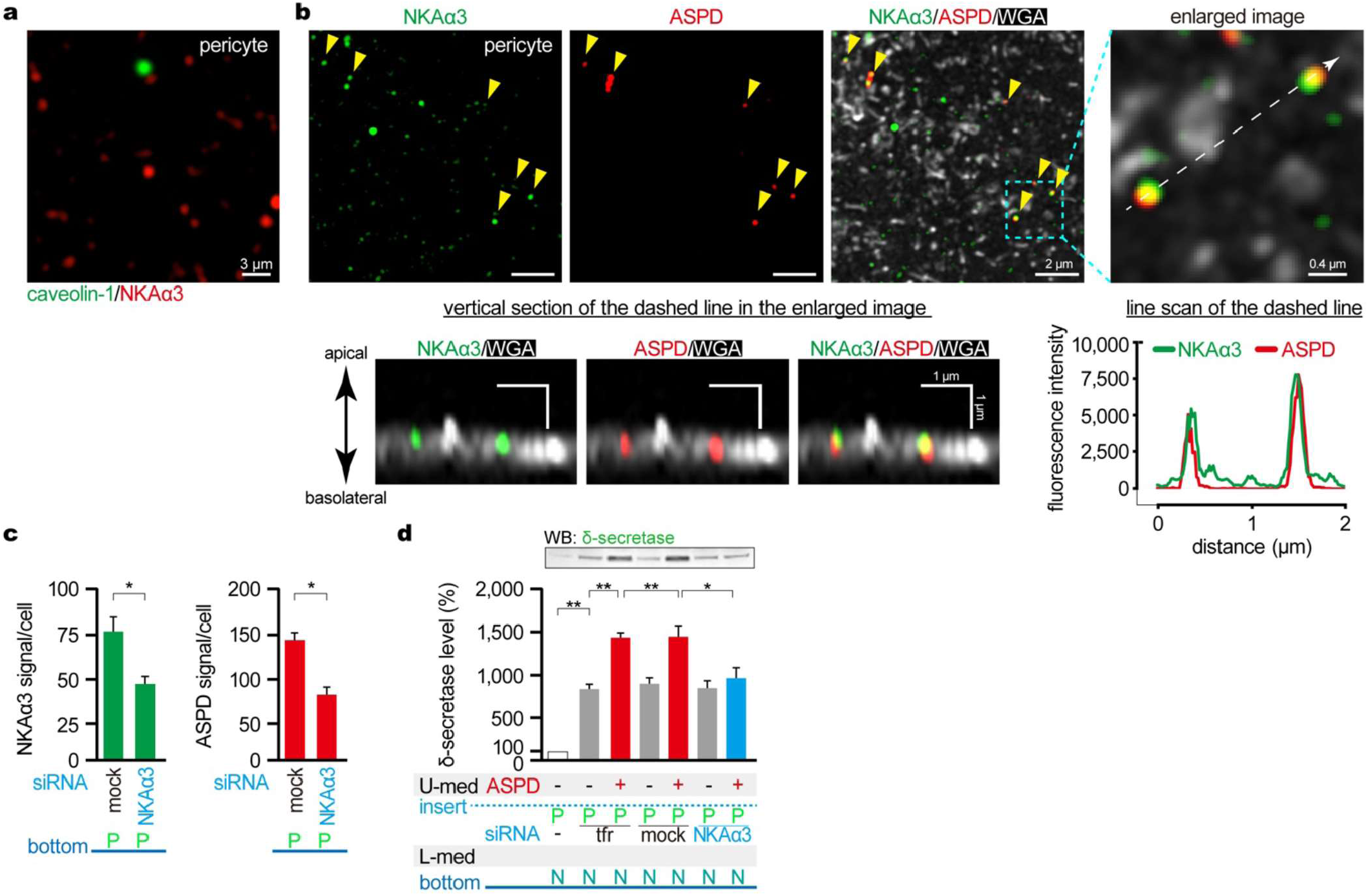
Identification of NKAα3 as an ASPD target in pericytes, leading to δ-secretase activation in neurons. **a**, Immunostaining (NKAα3, caveolin-1) of pericytes, showing lack of colocalization between NKAα3 and caveolin-1. **b**, Immunostaining (NKAα3, ASPD) of ASPD-treated (10 min, 38.9 nM) pericytes, showing the colocalization of ASPD and NKAα3 (yellow arrowheads) on the plasma membrane marked by fluorescent wheat germ agglutinin (WGA). The vertical sections and the line scan of the dashed line in the enlarged image are shown in the lower panels. **c**, Percentage of NKAα3 signals (*n* = 4) and ASPD signals (*n* = 8) following NKAα3 siRNA (30 nM) pretreatment of pericytes. NKAα3 or ASPD signals were counted by the immunostaining of pericytes treated with ASPD (10 min, 31.5 nM). **d**, Inhibition of δ-secretase activation in neurons by NKAα3 siRNA (30 nM) pretreatment of pericytes. siRNA (mock or NKAα3) was transfected to pericytes using the transfection reagent (tfr). Pericytes treated with siRNA or tfr alone were treated with ASPD (48 h, 41.1 nM), and δ-secretase levels were determined (*n* = 4) as in Fig. 1c. In **c-d**, data represent the mean ± S.E. \**p* < 0.05, \*\**p* < 0.01 (unpaired Welch’s t-test in **c** or one-way ANOVA with Scheffé’s method in **d**). For the raw data of **c** and **d**, refer to Extended Data Fig. 6**d, e** and **f**, respectively.

We then examined whether ASPD binds to NKAα3 in pericytes. Super-resolution microscopy showed that 85.5 ± 4.6% (*n* = 3) of ASPD interacted with NKAα3 10 min after ASPD treatment (Fig. 3b upper panels). The vertical section of the pericytes showed that ASPD and NKAα3 colocalized on the plasma membrane (Fig. 3b lower left). The line scan ensured that the fluorescence signals of ASPD and NKAα3 completely overlapped (Fig. 3b lower right). Moreover, knocking down pericyte NKAα3 by ∼40% with siRNA (Fig. 3c left and Extended Data Fig. 6c) also decreased the number of ASPD bound to pericytes by ∼40% (Fig. 3c right). Thus, in pericytes, the binding target of ASPD is also NKAα3.

Next, we studied whether pericyte NKAα3 mediates ASPD action leading to the activation of δ-secretase in neurons by knocking down pericyte NKAα3. Unexpectedly, we observed that the application of a transfection reagent alone to pericytes resulted in ∼9-fold activation of δ-secretase in neurons (Fig. 3d). Nevertheless, the administration of ASPD in addition to the transfection reagent produced ∼15-fold activation, 6-fold higher than the transfection reagent alone. siRNA treatment against NKAα3 eliminated this ASPD-dependent activation of δ-secretase (Fig. 3d). As pericytes and neurons do not contact directly, we postulate that an effector molecule, which is unknown at present, is released from pericytes upon ASPD binding to NKAα3 (but to some extent by the transfection reagent also) and activates δ-secretase in neurons.

Recent studies have suggested that neuronal δ-secretase may play a role in AD pathophysiology by cleaving APP and tau^25,33^. However, it remains largely unknown what pathological conditions or stimuli activate δ-secretase. We here found one such instance: APP_1-373,_ selectively produced by δ-secretase^25^, increased (Fig. 4a left); also, tau_1-368_, the δ-secretase-selective fragment of tau^33^, accumulated in neurons, whereas full-length tau decreased nearly 50% (Fig. 4a right and middle). Tau_1-368_ shows a reduced ability to promote tubulin polymerization into microtubules and is susceptible to hyperphosphorylation^33,34^.

**Fig. 4.**
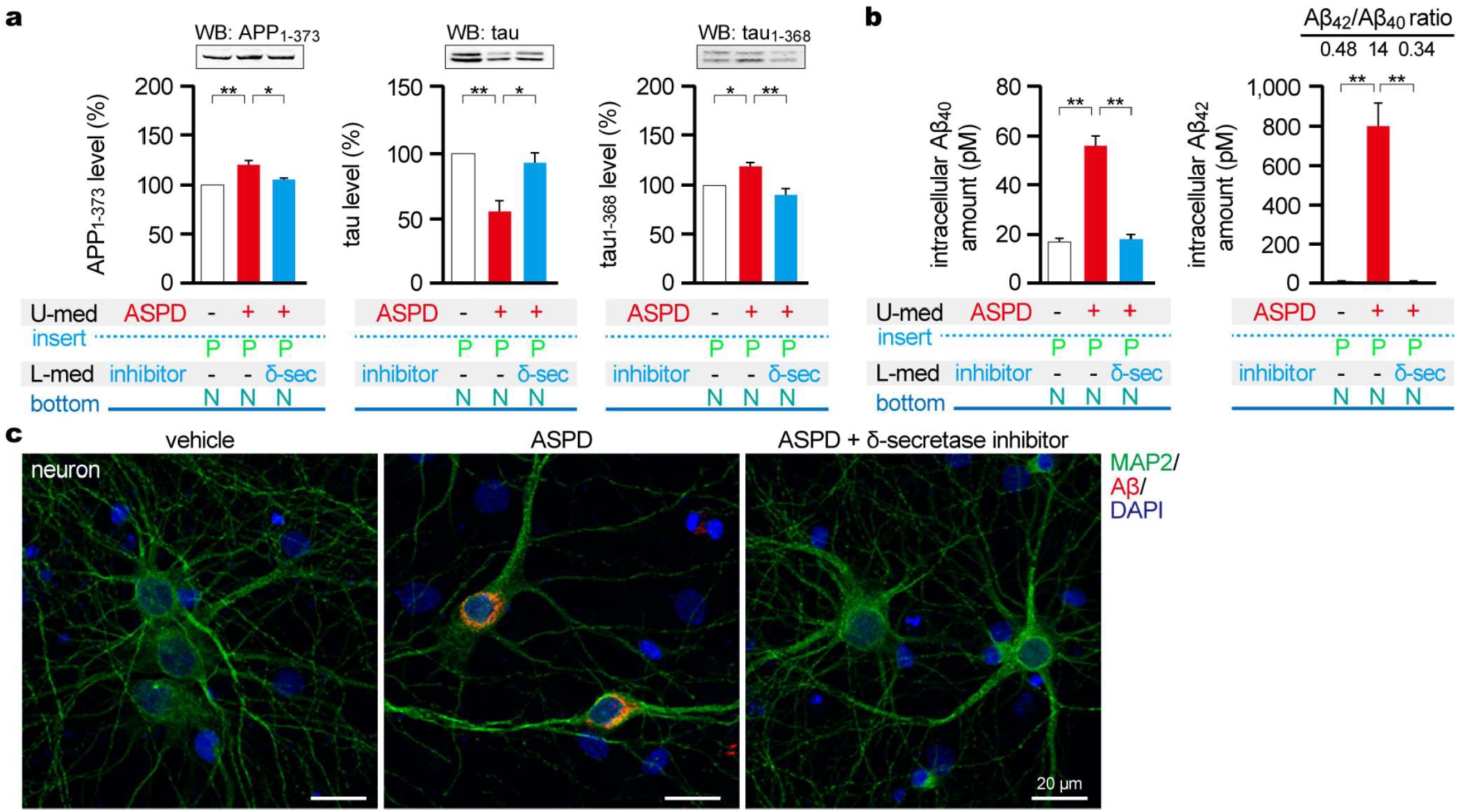
Fragmentation of APP and tau by δ-secretase induced by ASPD action on pericytes, leading to accumulation of Aβ and pathogenic tau_1-368_ fragment in neurons. **a**, Quantification by western blotting, normalized to actin (*n* = 3), of full-length tau and δ-secretase-specific tau_1-368_ fragment (double band due to the N-terminal insertion and multiple repeat regions in the microtubule-binding domain), and δ-secretase-specific APP_1-373_ fragment in neurons, with or without ASPD treatment (48 h, 43.4 nM) of pericytes. Treatment of neurons with a δ-secretase inhibitor (10 µM compound 11) blocked the ASPD-induced changes. **b and c**, Aβ increased in neurons expressing human APP Swedish mutation, as detected by Aβ ELISA (**b**) or immunostaining (**c**), after 48 h ASPD treatment (**b**, 43.4 nM; **c**, 60.0 nM) of pericytes. In **b**, the levels of Aβ_40_ and Aβ_42_ in the neuronal extracts were determined by ELISA specific to each Aβ (*n* = 3). In **c**, Aβ was detected by immunostaining with an N-terminal-end-specific antibody recognizing human Aβ but not APP. Treatment of neurons with the δ-secretase inhibitor blocked the Aβ increase in **b** and **c**. In **a** and **b**, data represent the mean ± S.E. \**p* < 0.05, \*\**p* < 0.01 (one-way ANOVA with Scheffé’s method). For the raw data of **a** and **b**, refer to Extended Data Fig. 7**a** and **b**, respectively.

Phosphorylated tau_1-368_ forms abnormal aggregates, eventually leading to neurofibrillary tangles (NFT)^33,35^, an AD marker, along with senile plaques consisting of abnormal Aβ aggregates. A δ-secretase-specific inhibitor (10 µM compound 11)^36^ abolished the increases of APP_1-373_ and tau_1-368_, indicating that δ-secretase is responsible for these changes (Fig. 4a).

Removal of the APP N-terminal region by δ-secretase facilitates subsequent cleavage by BACE1, resulting in doubled Aβ production^25^ (see also Fig. 1b). ASPD treatment of pericytes increased Aβ_40_ ∼3 times and Aβ_42_ as large as ∼130 times in neurons after 48 h of treatment, nearly twice the effect induced by the ASPD treatment of endothelial cells (Fig. 4b). The increases in Aβ_40_ and Aβ_42_ resulted in the accumulation of Aβ in the cell body of neurons expressing human APP Swedish mutation (Fig. 4c). The δ-secretase-specific inhibitor suppressed the increases as expected (Fig. 4b and c). Of note, ASPD treatment of endothelial cells or pericytes predominantly increased Aβ_42_, giving the Aβ_42_/Aβ_40_ ratio of 13-14, which is 0.1-0.2 in normal neurons^37^. The increased Aβ_42_/Aβ_40_ ratio in the brain indicates a stronger tendency for aggregation, reflecting that Aβ_42_ is more prone to aggregate than Aβ_40_ and is associated with AD development in both sporadic and familial cases^37^.

It is documented that, upon stimulation, pericytes secrete effector molecules upon stimulation, which influence and/or control surrounding endothelial cells and astrocytes^38,39^. Our results suggest that pericytes control metabolism in neurons through effector molecules, one of which ASPD increases abnormally, and that too high a concentration of the effector molecule in the extracellular medium activates δ-secretase in healthy neurons.

### ASPD, BACE1 and δ-secretase are linked in AD

Using our NVU mimicking *in-vitro* 3D culture system, we have so far demonstrated that ASPD in blood imposes Aβ and tau pathologies on healthy neurons by increasing BACE1 and δ-secretase (Fig. 6). We extended the analysis to patient brains (from five people in a preclinical state and five patients suffering from AD; see Extended Data Table 1 for details). The levels of BACE1, δ-secretase and δ-secretase-specific fragments (APP_1-373_ and tau_1-368_) in autopsy brains (dissected within 4.7 ± 0.7 h of death) were determined by western blotting. In parallel, the amount of ASPD was quantified by dot blotting using an ASPD-specific antibody^7^. Compared to the preclinical cases, ASPD, BACE1, δ-secretase, APP_1-373_ and tau_1-368_ all increased significantly in the AD cases (Fig. 5 upper panels). Notably, the ASPD amount positively correlated with the levels of BACE1, δ-secretase, APP_1-373_ and tau_1-368_ (Fig. 5 lower panels). BACE1 and δ-secretase, belonging to two different signaling cascades, also showed a positive correlation. The positive correlations support the view that ASPD increases BACE1 and δ-secretase in patient brains, although it is necessary to quantify them in a larger number of autopsy brains.

**Fig. 5.**
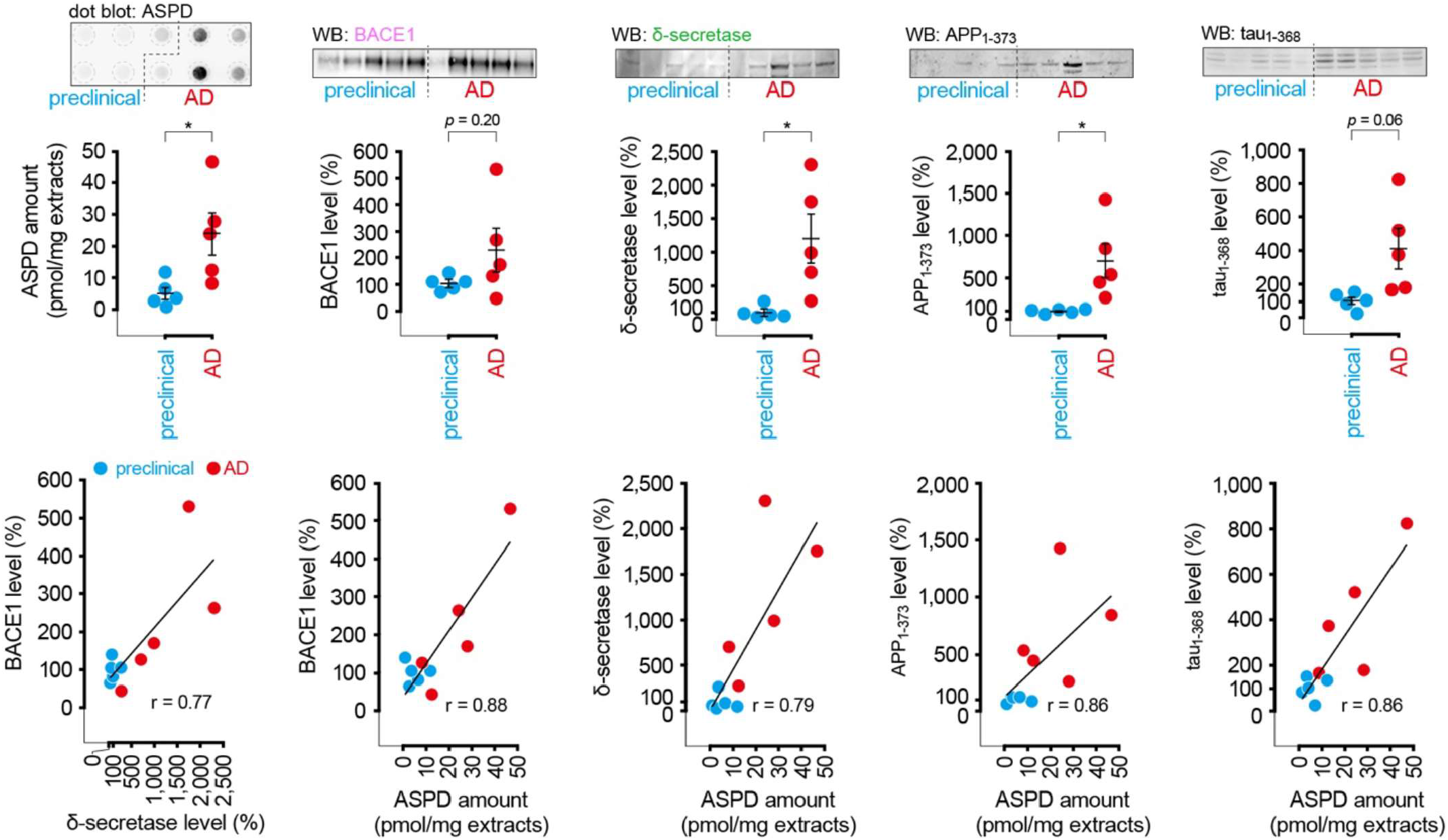
Correlation of ASPD amount with protein levels of BACE1, active δ-secretase, APP_1-373_ and tau_1-368_ in preclinical and AD patient brains. Frozen brain blocks were obtained from the occipital cortex of freshly frozen autopsy brains derived from five individuals in the preclinical state (Braak stage II/III for neurofibrillary tangles (NFT) and CERAD score A-C for senile plaques) and from five individuals with AD (Braak stage V/VI for NFT and CERAD score C for senile plaques; details in Extended Data Table 1). Total proteins were extracted from half of the blocks with RIPA buffer and used for the quantification by western blotting (*n* = 3) of BACE1, δ-secretase, APP_1-373_ and tau_1-368_ (triple band due to variations in the N-terminal insertion and multiple repeat regions in the microtubule-binding domain) (Extended Data Fig. 8). Soluble proteins were extracted^7^ from the other half of the blocks and used for the purification of ASPD^7^. ASPD was quantified by dot blotting (1 µg/dot) with ASPD-specific rpASD1 antibody^7^ (*n* = 3). As for the levels of BACE1, active δ-secretase, APP_1-373_ and tau_1-368_, the average values in the preclinical group were taken as 100%. Data represent the mean ± S.E. normalized to actin levels. \**p* < 0.05 (the unpaired Welch’s *t*-test). Lower panels show the correlation analysis. Pearson’s correlation values (r) were calculated using all values (*n* = 10). For the raw data of the upper panels, refer to Extended Data Fig. 8.

**Fig. 6.**
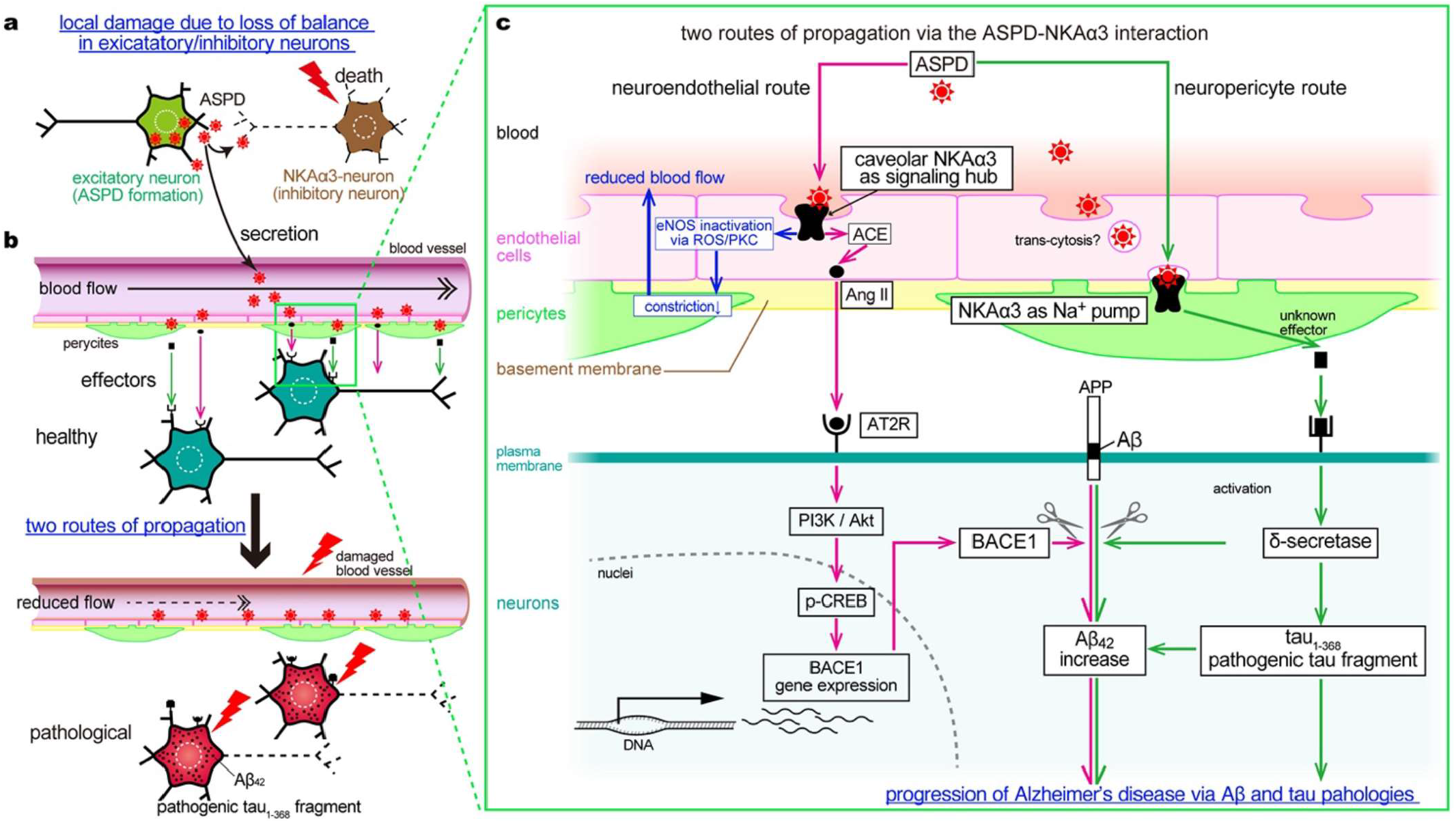
Multiple roles of ASPD via NKAα3 in pathogenesis and progression of AD. **a**, Local neuronal damage induced by ASPD-NKAα3. ASPD is formed in excitatory neurons and secreted, leading to the degeneration of neighboring NKAα3-expressing inhibitory neurons via the inhibition of NKAα3 pump activity^8,14^. This causes hyperexcitation of the excitatory neurons and the production of more Aβ, resulting in ASPD accumulation. This series of events, presumably an electrical feedback mechanism, destroys the balance of excitatory/inhibitory neurons and eventually kills both neurons. Such excitation/inhibition imbalance is the mechanism that works locally but also between cortical layers. **b** and **c**, Two routes of propagation of the pathology induced by ASPD-NKAα3.

## Discussion

AD progresses over many years and through multiple distinct stages. It begins with the accumulation of Aβ aggregates in a limited cortical region. The secretion of fragmented tau follows, then its phosphorylation. Subsequently, tau aggregates (NFT) accumulate in the cerebral cortex. Eventually, widespread neuronal loss and cognitive impairment take place. This order of events is strict, and the stage in AD can now be determined quantitatively, as demonstrated by a recent study on cerebrospinal fluid (CSF) biomarkers using two large cohorts^3^. One of the outstanding features of AD is that tau pathology supersedes Aβ pathology. As the gene groups related to Aβ and tau pathologies show no overlap^19^, the underlying molecular mechanisms have been thought to be different. Yet, as every AD patient follows the identical course^3^, there must be a mechanism linking them.

Another outstanding feature in the early stage of AD is the rapid spread of Aβ pathology. The aggregation of Aβ occurs inside neurons in the early stage of AD, with ASPD, ∼30-mer of Aβ, as the main component^14,16^. Only excitatory neurons produce and secrete ASPD^14^, but secreted ASPD impairs inhibitory neurons projecting to the excitatory neurons, by binding to NKAα3 (Fig. 6a). The impairment of inhibitory neurons causes the hyperexcitation of excitatory neurons and the production of more Aβ, resulting in ASPD accumulation. This series of events, presumably an electrical feedback mechanism, destroys the balance of excitatory/inhibitory neurons and eventually kills both neurons (Fig. 6a)^14,16^. This loss of balance is a mechanism that works locally but also between cortical layers, as the loss of balance in one layer leads to an imbalance in another layer that connects to it due to the column structure running vertically through the cortical layers. However, such a mechanism does not explain the rapid spread of Aβ pathology to far distant brain regions not directly connected. This rapid spread could be explained if a signal or a signaling cascade works through the environment surrounding neurons (e.g., blood or extracellular fluids). Since every neuron produces small amounts of Aβ for synaptic connection, it is only necessary to transfer conditions (effector molecules) that urge Aβ production and aggregation. In this study, we have identified two such signaling cascades that work through endothelial cells and pericytes in capillaries (Fig. 6b and c). Moreover, one of the cascades triggers tau pathology. At the origin of either cascade, we identified ASPD and its sole target, NKAα3, generally described as a neuron-specific isoform of Na^+^,K^+^-ATPase but present in endothelial cells and even abundant in pericytes.

These two cascades are distinct in several aspects. In endothelial cells, ASPD only acts on caveolar NKAα3 and produces Ang II through ACE. The released Ang II induces BACE1 via the AT2R/PI3K/Akt/p-CREB signaling cascade, causing a huge increase of Aβ_42_ and presumably its aggregates, as Aβ_42_ forms aggregates much more easily than Aβ_40_. This signaling cascade is independent of the ROS/PKC/eNOS signaling cascade that we previously identified in endothelial cells^22^. The ROS/PKC/eNOS signaling cascade reduces NO released from endothelial cells, inhibits capillary contraction, and reduces blood flow (Fig. 6b). Because the Ang II and eNOS cascades are independent, caveolar NKAα3 appears to act as a signaling hub, as postulated for NKAα1^40^ (Fig. 6c).

In pericytes, ASPD binds to non-caveolar NKAα3 and activates a totally different signaling cascade. In marked contrast to endothelial cells, pericytes are abundant with NKAα3 but almost devoid of caveolin-1. The unexpected abundance of NKAα3 suggests that α3 is the isoform suitable for excitable cells, as pericytes are excitable, like neurons. ASPD-binding to NKAα3 in pericytes releases an effector molecule that is currently unknown. This process activates δ-secretase, which increases pathological Aβ_42_ further but also produces pathogenic tau_1-368_ fragment, observations confirmed with human autopsy brains. Previous studies on cohorts^35,41^ and disease model mice (5xFAD and TauP301S mice)^36^ support the activation of δ-secretase in AD: tau_368_, enriched in NFT, can be a CSF biomarker that reflects cognitive impairment and NFT better than CSF phosphorylated tau^35,41^. The δ-secretase inhibitor, also used in this study, reduces the δ-secretase-driven degradation of APP and tau and attenuates learning impairment in the model mice^36^. Nevertheless, the actual role of δ-secretase in the onset of AD was ambiguous because when and how δ-secretase is activated was unknown. We now see that δ-secretase is pivotal in linking Aβ and tau pathologies (Fig. 6c).

ACE is another molecule, the role of which in AD has been questioned, even though its increase in AD brains has been repeatedly reported^42,43^. This is because ACE shows an ability to degrade Aβ in vitro^44^, suggesting its contribution to a biological defense reaction. Our results clearly show that ACE plays a role in spreading Aβ pathology, consistent with a recent unbiased analysis of two large cohorts^19^. That study showed ACE expression is higher in the brain areas more susceptible to Aβ accumulation and that an increase in ACE is inversely correlated with the score of the Montreal cognitive assessment, a test for detecting mild cognitive impairment^19^. Accordingly, it is reasonable to expect that when ASPD in the blood exceeds a certain amount, effector molecules, including Ang II, that promote Aβ pathology are released from the capillaries. Consequently, Aβ aggregates begin to accumulate simultaneously in wide and distant regions.

In this study, we have elucidated a part of the scenario for the stage of AD in which Aβ pathology spreads and secretory tau pathology begins. The underlying mechanism is the control through the NVU comprising blood capillary and pericytes with effector molecules. As it spreads throughout the brain, the capillary network can be a useful means of synchronizing or imposing the same “conditions” over large, possibly not directly connected brain regions. Conversely, if pathological conditions occur, the NVU may work as a machinery for rapidly spreading the disease to a wide area of the brain. Thus, stopping the signal cascades that spread Aβ pathology and activate tau pathology near the origin may halt the progression of AD. Because ASPD and NKAα3 work at the origins of the cascades, drugs that inhibit the binding of ASPD to NKAα3, as we previously discovered^8^, may suppress the onset and progression of AD in both neurons and blood vessels.

## Acknowledgments

This work is supported by a Grants-in-Aid for Young Scientists (20K16022 to T.S.), and for Scientific Research (22K06643 to T.S.; 20H03457 to M.H.) from Japan Society for the Promotion of Science; a Collaborative Research Project of the Brain Research Institute, Niigata University to M.H.; a Medical Research Grant (to T.S.) and Specific Research Grants (to M.H) from Takeda Science Foundation; and a grant from AMED (No. JP22wm0525037 to M.H.; JP23wm0425019 to A.K.). The authors thank Dr. David B. Teplow (University of California, Los Angeles) and Dr. Chikashi Toyoshima (The University of Tokyo) for discussions.

## Author contributions

T.S. and M.H. designed the study, wrote and edited the manuscript and the prepared figures; T.S. generated the 3D culture system, performed the experiments and analyzed the data; M.H. analyzed the data, provided feedback and supervised all research; S.H. and A.K. selected and prepared the autopsy brains; M.I., N.T. and S.M. produced an AAV vector carrying human APP Swedish mutation; all authors read, edited and approved the final manuscript.

## Competing interest declaration

M.H. has served as a technical advisor to TAO Health Life Pharma Co. Ltd., a Kyoto University-derived bio venture, with the permission of the conflict-of-interest committee of FBRI. T.S. was an employee of TAO Health Life Pharma Co. Ltd. until April 2019.

## Notes

### Competing Interest Statement

Minako Hoshi has served as a technical advisor to TAO Health Life Pharma Co. Ltd., a Kyoto University-derived bio venture, with the permission of the conflict-of-interest committee of FBRI. Tomoya Sasahara was an employee of TAO Health Life Pharma Co. Ltd. until April 2019.

## Main references

1 Selkoe, D. J. & Hardy, J. The amyloid hypothesis of Alzheimer’s disease at 25 years. EMBO Mol Med 8, 595–608 (2016). 10.15252/emmm.201606210

2 Hoshi, M. Tracking down a missing trigger for Alzheimer’s disease by mass spectrometric imaging based on brain network analysis. Prog Mol Biol Transl Sci 168, 25–55 (2019). 10.1016/bs.pmbts.2019.05.011

3 Salvado, G. et al. Disease staging of Alzheimer’s disease using a CSF-based biomarker model. Nat Aging (2024). 10.1038/s43587-024-00599-y

4 Hamilton, N. B., Attwell, D. & Hall, C. N. Pericyte-mediated regulation of capillary diameter: a component of neurovascular coupling in health and disease. Front Neuroenergetics 2 (2010). 10.3389/fnene.2010.00005

5 Merlini, M., Davalos, D. & Akassoglou, K. In vivo imaging of the neurovascular unit in CNS disease. Intravital 1, 87–94 (2012). 10.4161/intv.22214

6 Hoshi, M. et al. Spherical aggregates of β-amyloid (amylospheroid) show high neurotoxicity and activate tau protein kinase I/glycogen synthase kinase-3β. Proc. Natl. Acad. Sci. U.S.A. 100, 6370–6375 (2003). 10.1073/pnas.1237107100

7 Noguchi, A. et al. Isolation and characterization of patient-derived, toxic, high mass amyloid β-protein (Aβ) assembly from Alzheimer disease brains. J Biol Chem 284, 32895–32905 (2009). 10.1074/jbc.M109.000208

8 Ohnishi, T. et al. Na, K-ATPase α3 is a death target of Alzheimer patient amyloid-β assembly. Proc Natl Acad Sci U S A 112, E4465–4474 (2015). 10.1073/pnas.1421182112

9 Blanco, G. & Mercer, R. W. Isozymes of the Na-K-ATPase: heterogeneity in structure, diversity in function. Am J Physiol 275, F633–650 (1998). 10.1152/ajprenal.1998.275.5.F633

10 McGrail, K. M., Phillips, J. M. & Sweadner, K. J. Immunofluorescent localization of three Na,K-ATPase isozymes in the rat central nervous system: both neurons and glia can express more than one Na,K-ATPase. J Neurosci 11, 381–391 (1991). 10.1523/JNEUROSCI.11-02-00381.1991

11 Haass, C., Kaether, C., Thinakaran, G. & Sisodia, S. Trafficking and proteolytic processing of APP. Cold Spring Harb Perspect Med 2, a006270 (2012). 10.1101/cshperspect.a006270

12 Cai, W., Li, L., Sang, S., Pan, X. & Zhong, C. Physiological Roles of β-amyloid in Regulating Synaptic Function: Implications for AD Pathophysiology. Neurosci Bull 39, 1289–1308 (2023). 10.1007/s12264-022-00985-9

13 Benilova, I., Karran, E. & De Strooper, B. The toxic Aβ oligomer and Alzheimer’s disease: an emperor in need of clothes. Nat Neurosci 15, 349–357 (2012). 10.1038/nn.3028

14 Komura, H. et al. Alzheimer Aβ Assemblies Accumulate in Excitatory Neurons upon Proteasome Inhibition and Kill Nearby NAKα3 Neurons by Secretion. iScience 13, 452–477 (2019). 10.1016/j.isci.2019.01.018

15 Murata, K. et al. Region- and neuronal-subtype-specific expression of Na,K-ATPase alpha and beta subunit isoforms in the mouse brain. J Comp Neurol (2020). 10.1002/cne.24924

16 Hoshi, M. Multi-angle development of therapeutic methods for Alzheimer’s disease. Br J Pharmacol 178, 770–783 (2021). 10.1111/bph.15174

17 Yu, M., Sporns, O. & Saykin, A. J. The human connectome in Alzheimer disease - relationship to biomarkers and genetics. Nat Rev Neurol 17, 545–563 (2021). 10.1038/s41582-021-00529-1

18 Buckner, R. L., Andrews-Hanna, J. R. & Schacter, D. L. The brain’s default network: anatomy, function, and relevance to disease. Ann N Y Acad Sci 1124, 1–38 (2008). 10.1196/annals.1440.011

19 Yu, M. et al. Spatial transcriptomic patterns underlying amyloid-β and tau pathology are associated with cognitive dysfunction in Alzheimer’s disease. Cell Rep 43, 113691 (2024). 10.1016/j.celrep.2024.113691

20 Andjelkovic, A. V., Stamatovic, S. M., Phillips, C. M., Martinez-Revollar, G. & Keep, R. F. Modeling blood-brain barrier pathology in cerebrovascular disease in vitro: current and future paradigms. Fluids Barriers CNS 17, 44 (2020). 10.1186/s12987-020-00202-7

21 Snowdon, D. A. et al. Brain infarction and the clinical expression of Alzheimer disease. The Nun Study. JAMA 277, 813–817 (1997).

22 Sasahara, T., Satomura, K., Tada, M., Kakita, A. & Hoshi, M. Alzheimer’s Aβ assembly binds sodium pump and blocks endothelial NOS activity via ROS-PKC pathway in brain vascular endothelial cells. iScience 24, 102936 (2021). 10.1016/j.isci.2021.102936

23 Kang, S. S., Ahn, E. H. & Ye, K. Delta-secretase cleavage of Tau mediates its pathology and propagation in Alzheimer’s disease. Exp Mol Med 52, 1275–1287 (2020). 10.1038/s12276-020-00494-7

24 Vassar, R. et al. β-secretase cleavage of Alzheimer’s amyloid precursor protein by the transmembrane aspartic protease BACE. Science 286, 735–741 (1999). 10.1126/science.286.5440.735

25 Zhang, Z. et al. Delta-secretase cleaves amyloid precursor protein and regulates the pathogenesis in Alzheimer’s disease. Nat Commun 6, 8762 (2015). 10.1038/ncomms9762

26 Timmermans, P. B. & Smith, R. D. Angiotensin II receptor subtypes: selective antagonists and functional correlates. Eur Heart J 15 Suppl D, 79–87 (1994). 10.1093/eurheartj/15.suppl_d.79

27 Nilsberth, C. et al. The ‘Arctic’ APP mutation (E693G) causes Alzheimer’s disease by enhanced Aβ protofibril formation. Nat Neurosci 4, 887–893 (2001). 10.1038/nn0901-887

28 Ancolio, K. et al. Unusual phenotypic alteration of β amyloid precursor protein (βAPP) maturation by a new Val-715 → Met βAPP-770 mutation responsible for probable early-onset Alzheimer’s disease. Proc Natl Acad Sci U S A 96, 4119–4124 (1999). 10.1073/pnas.96.7.4119

29 Choi, G. E. et al. Membrane-Associated Effects of Glucocorticoid on BACE1 Upregulation and Aβ Generation: Involvement of Lipid Raft-Mediated CREB Activation. J Neurosci 37, 8459–8476 (2017). 10.1523/JNEUROSCI.0074-17.2017

30 Xie, F. et al. Identification of a Potent Inhibitor of CREB-Mediated Gene Transcription with Efficacious in Vivo Anticancer Activity. J Med Chem 58, 5075–5087 (2015). 10.1021/acs.jmedchem.5b00468

31 Hashikawa-Hobara, N. et al. Candesartan cilexetil improves angiotensin II type 2 receptor-mediated neurite outgrowth via the PI3K-Akt pathway in fructose-induced insulin-resistant rats. Diabetes 61, 925–932 (2012). 10.2337/db11-1468

32 Johannessen, M., Delghandi, M. P. & Moens, U. What turns CREB on? Cell Signal 16, 1211–1227 (2004). 10.1016/j.cellsig.2004.05.001

33 Zhang, Z. et al. Cleavage of tau by asparagine endopeptidase mediates the neurofibrillary pathology in Alzheimer’s disease. Nat Med 20, 1254–1262 (2014). 10.1038/nm.3700

34 Hu, W., Tung, Y. C., Zhang, Y., Liu, F. & Iqbal, K. Involvement of Activation of Asparaginyl Endopeptidase in Tau Hyperphosphorylation in Repetitive Mild Traumatic Brain Injury. J Alzheimers Dis 64, 709–722 (2018). 10.3233/JAD-180177

35 Blennow, K. et al. Cerebrospinal fluid tau fragment correlates with tau PET: a candidate biomarker for tangle pathology. Brain 143, 650–660 (2020). 10.1093/brain/awz346

36 Zhang, Z. et al. Inhibition of delta-secretase improves cognitive functions in mouse models of Alzheimer’s disease. Nat Commun 8, 14740 (2017). 10.1038/ncomms14740

37 Aoki, M., Volkmann, I., Tjernberg, L. O., Winblad, B. & Bogdanovic, N. Amyloid β-peptide levels in laser capture microdissected cornu ammonis 1 pyramidal neurons of Alzheimer’s brain. Neuroreport 19, 1085–1089 (2008). 10.1097/WNR.0b013e328302c858

38 Armulik, A., Genove, G. & Betsholtz, C. Pericytes: developmental, physiological, and pathological perspectives, problems, and promises. Dev Cell 21, 193–215 (2011). 10.1016/j.devcel.2011.07.001

39 van Splunder, H., Villacampa, P., Martinez-Romero, A. & Graupera, M. Pericytes in the disease spotlight. Trends Cell Biol 34, 58–71 (2024). 10.1016/j.tcb.2023.06.001

40 Tian, J. et al. Binding of Src to Na+/K+-ATPase forms a functional signaling complex. Mol Biol Cell 17, 317–326 (2006). 10.1091/mbc.e05-08-0735

41 Simren, J. et al. CSF tau368/total-tau ratio reflects cognitive performance and neocortical tau better compared to p-tau181 and p-tau217 in cognitively impaired individuals. Alzheimers Res Ther 14, 192 (2022). 10.1186/s13195-022-01142-0

42 Miners, J. S. et al. Angiotensin-converting enzyme (ACE) levels and activity in Alzheimer’s disease, and relationship of perivascular ACE-1 to cerebral amyloid angiopathy. Neuropathol Appl Neurobiol 34, 181–193 (2008). 10.1111/j.1365-2990.2007.00885.x

43 Miners, S. et al. Angiotensin-converting enzyme levels and activity in Alzheimer’s disease: differences in brain and CSF ACE and association with ACE1 genotypes. Am J Transl Res 1, 163–177 (2009).

44 Hu, J., Igarashi, A., Kamata, M. & Nakagawa, H. Angiotensin-converting enzyme degrades Alzheimer amyloid β-peptide (Aβ); retards Aβ aggregation, deposition, fibril formation; and inhibits cytotoxicity. J Biol Chem 276, 47863–47868 (2001). 10.1074/jbc.M104068200

